# MiR-29b-1-5p is altered in BRCA1 mutant tumours and is a biomarker in basal-like breast cancer

**DOI:** 10.1101/319129

**Authors:** Michael J.G. Milevskiy, Gurveen K. Sandhu, Anna Wronski, Darren Korbie, Brooke L. Brewster, Annette Shewan, Stacey L. Edwards, Juliet D. French, Melissa A. Brown

**Affiliations:** School of Chemistry and Molecular Biosciences, University of Queensland, St Lucia, Queensland, Australia; Australian Institute of Biotechnology and Nanotechnology, University of Queensland, St Lucia, Queensland, Australia; QIMR Berghofer Medical Research Institute, Brisbane, Queensland, Australia

## Abstract

Depletion of BRCA1 protein in mouse mammary glands results in defects in lactational development and increased susceptibility to mammary cancer. Extensive work has focussed on the role of BRCA1 in the normal breast and in the development of breast cancer, the cell of origin for BRCA1 tumours and the protein-coding genes altered in BRCA1 deficient cells. However, the role of non-coding RNAs in BRCA1-deficient cells is poorly understood. To evaluate miRNA expression in BRCA1 deficient mammary cells, RNA sequencing was performed on the mammary glands of Brca1 knockout mice. We identified 140 differentially expressed miRNAs, 9 of which were also differentially expressed in human BRCA1 breast tumours or familial non-BRCA1 patients and during normal gland development. We show that BRCA1 binds to putative *cis*-elements in promoter regions of the miRNAs and regulates their expression, and that four miRNAs (miR-29b-1-5p, miR-664, miR-16-2 and miR-744) significantly stratified the overall survival of basal-like tumours. Importantly the prognostic value of miR-29b-1-5p was higher in significance than several commonly used clinical biomarkers. These results emphasize the role of Brca1 in modulating expression of miRNAs and highlights the potential for BRCA1 regulated miRNAs to be informative biomarkers associated with BRCA1 loss and survival in breast cancer.

## Introduction

Germline mutations in the breast cancer susceptibility gene, *BRCA1*, confer a high risk of developing neoplastic lesions. To understand how BRCA1-deficient cells give rise to BRCA1 breast tumours, extensive research has been performed to analyse and dissect the molecular expression profiles, cell of origin and genetic pathways linked to BRCA1 ^1−5^. BRCA1-deficient breast tumours often present as difficult to treat triple negative breast cancers (TNBC) not dissimilar to the basal-like molecular subtype of breast cancer, which lack expression of hormone receptors and easy to target growth signals. Whilst significant discoveries have been made on the contribution of the coding genome, the role of non-coding RNAs, such as microRNAs (miRNAs), in BRCA1-associated tumourigenesis remains unclear.

MiRNAs are small non-coding RNA molecules that predominantly inhibit gene expression, post-transcriptionally ^6^. MiRNAs function through two distinct mechanisms, perfect complementary binding or imperfect binding to mRNA 3’ untranslated regions (UTRs) leading to mRNA degradation or inhibition of translation, respectively. MiRNAs have key roles in tumorigenesis and development of breast cancer with widespread affects across all hallmarks of cancer ^7,8^.

BRCA1 is essential for DNA repair via homologous recombination, but is also a direct effector of gene expression. Recent ChIP-Seq data has shown that BRCA1 binds to promoters of numerous genes in human breast cell lines mediating downstream effects of NF-κB, TNF-α and retinoic acid (RA) growth signals ^9^. RA growth suppression was further reduced in BRCA1-knockdown cells and the mutant cell line HCC1937, which expresses a truncated protein incapable of binding to DNA ^9^. BRCA1 has also been shown to bind to and regulate miR-155 expression, in breast cancer ^10^. A BRCA1 R1699Q, a point mutation found in non-familial breast cancer patients, results in the failure of BRCA1 to recruit HDAC2 to the miR-155 promoter leading to histone acetylation and a subsequent increase in miR-155 expression contributing to breast tumorigenesis ^10^.

Knockout of Brca1 in murine mammary glands leads to defective development, which is most evident during lactation where aberrant growth of the lobular alveolar compartment is observed ^11,12^. Tumours arsing in Brca1 knockout mice share many morphological characteristics with their human counterparts including abnormal nuclei and high mitotic index ^13^. These tumours also share many fundamental characteristics including aberrant DNA damage, reminiscent of a defective Brca1 pathway, and gene expression signatures of basal-like breast tumours ^14,15^, therefore, we reasoned that represents a robust model for investigation of miRNAs dysregulation in Brca1-associated breast tumours.

We have previously identified a number of microRNAs deregulated following conditional loss of Brca1 in the mouse mammary gland ^16^. To further our studies, we performed microRNA-seq (miR-Seq) on Brca1 deficient murine mammary glands to evaluate the changes in miRNA expression. Our findings will facilitate a better understanding of basal-like and BRCA1-associated tumours and provide tools for greater patient prognostication.

## Results

### miRNAs are differentially expressed in *Brca1* knockout mammary glands

To identify miRNAs differentially expressed in Brca1 knockout mammary glands, miRNA-seq analysis was performed using RNA extracted from mammary glands at day 1 of lactation. 140 miRNAs were differentially expressed, of which 39 were down-regulated and 101 were up-regulated in MMTV-Cre/Brca1^fl/fl^ glands compared to Brca1^fl/fl^ controls (Figure 1A, Supp Table 1). Notably, five miRNAs, previously identified using a candidate approach, were differentially expressed (miR-205, miR-31, miR-148a, miR-181c, miR-200b and miR-210), confirming the validity of our approach.

**Figure 1.**
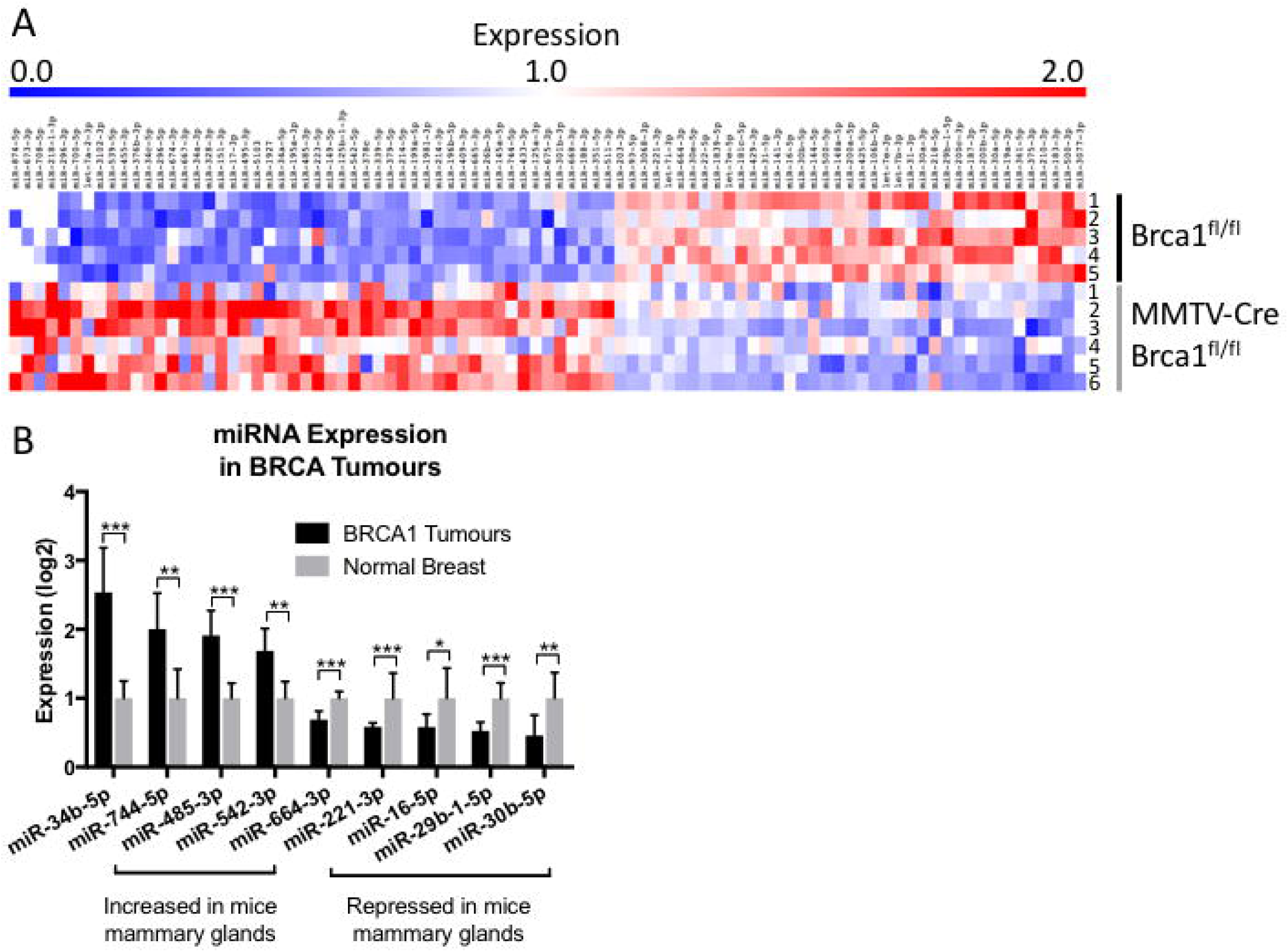
Brca1 deregulates miRNA expression in the mammary gland. **A**, summary of miRNA-seq data from mammary glands at day 1 of lactation in Brca1^fl/fl^ (n=5) and MMTV-Cre/Brca1^fl/fl^ glands (n=6), mouse number indicated on the right of heat map. **B**, expression of miRNAs with an identical trend of expression between murine *Brca1* deficient mammary tissue and human *BRCA1* mutant tumours. Nine out of 140 miRNAs were significantly conserved in expression between murine *Brca1* deficient tissue and human *BRCA1* mutant tumours. Statistical significance was measured by Students’ t-Test, p-values listed in Supp Table 1. (\**P≤0.05*, \*\**P≤0.01*, \*\*\**P≤0.001*).

### Differentially expressed miRNAs are deregulated in *BRCA1*-associated tumours

Mechanisms of transcriptional regulation can often be conserved between developmental processes and cancer. To explore this, we examined the expression of miRNAs differentially expressed in Brca1 deficient mammary glands with those miRNAs altered in 13 breast tumours of human BRCA1 mutation carriers (Published cohort ^17^). Nine miRNAs were identified, four which displayed elevated expression (miR-34b-5p, miR-744-p, miR-485-3p, miR-542-3p) in both *Brca1* knockout mammary glands and BRCA1 breast tumours and five were decreased (miR-664-3p, miR-221-3p, miR-16-5p, miR-29b-1-5p and miR-30b-5p) in both datasets (Figure 1B, Supp Table 2).

### Differentially expressed miRNAs are deregulated in the murine mammary gland

Brca1 is important for the development of the mammary gland, particularly for the maintenance of the luminal cell progenitor pool and proper gene expression during lactation ^1,12^. Using published Illumina expression array data, we evaluated five of the nine differentially expressed miRNAs present on the array in murine mammary glands from 12 days of age through pregnancy and the final stages of involution from 2-3 mice per age group. Each miRNA demonstrated differential expression throughout the developmental stages of the mammary gland (Figure 2A). Interestingly, four miRNAs (miR-34b-5p, miR-221-3p, miR-29b-1-5p and miR-30b-5p) display decreased expression at lactation day 5 compared to the final stages of gestation suggesting that loss of these miRNAs may be required for milk production and the final involution stage of the mammary gland. MiR-221 and miR-29b-1 positively correlated with Brca1 expression supporting the finding that these miRNAs decrease following Brca1 knockout in the mouse and human patients with BRCA1 mutations.

**Figure 2.**
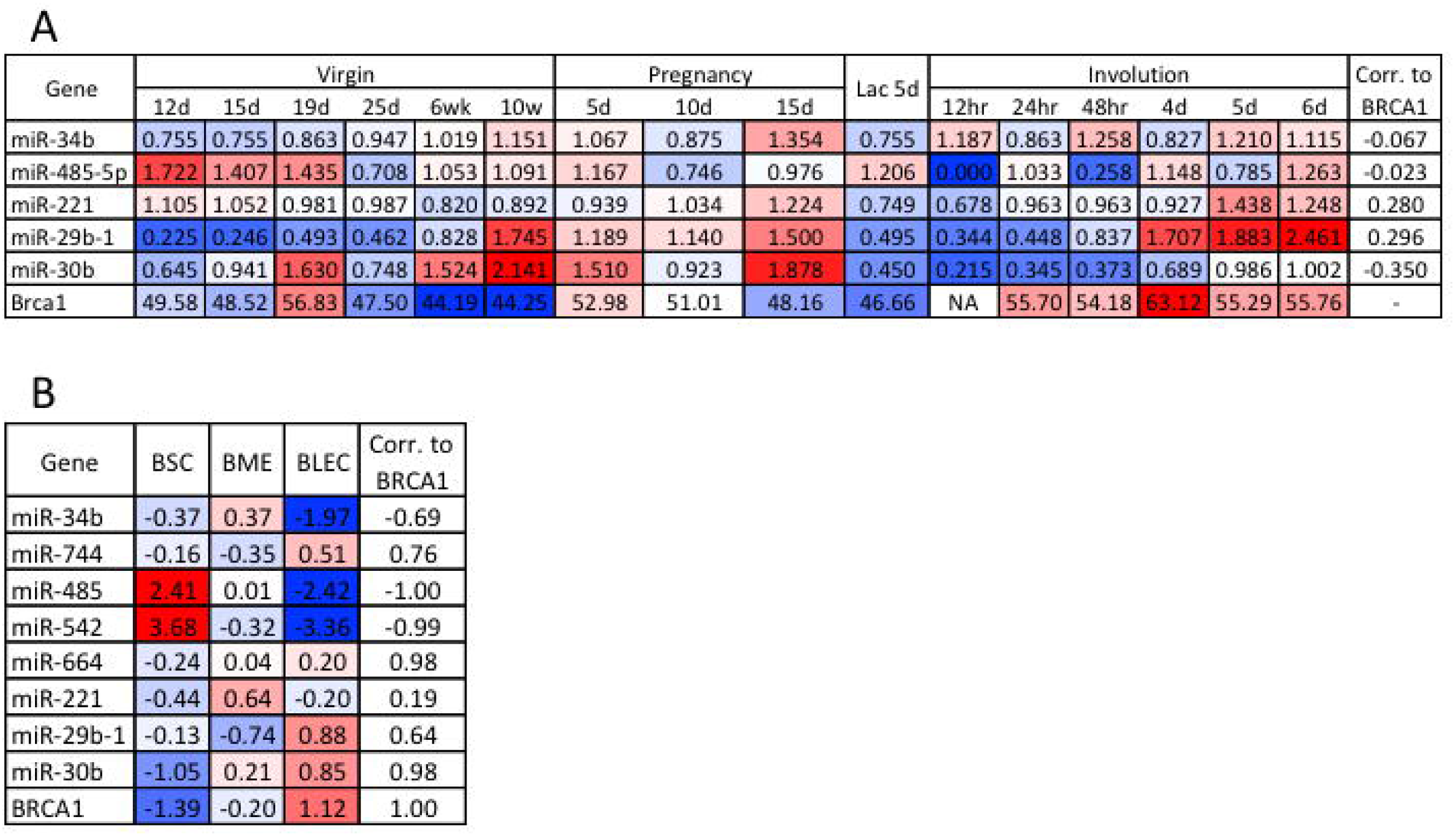
Deregulated miRNAs are differentially expressed in the mammary gland. **A,** miRNA and *Brca1* mRNA expression data from murine mammary glands through puberty, pregnancy, lactation and involution. Define hr, d, wk **B,** miRNA and *BRCA1* mRNA expression data from BSC (breast stem cell-like, n=2), BME (breast myoepithelial, n=2) and BLEC (breast luminal epithelial cell, n=2) epithelial populations of the human breast.

Utilising a published miRNA-Seq dataset we next interrogated expression in human mammary organoids comprised of myoepithelial, luminal and stem-like cells, which strongly resemble their *in vivo* counterparts that comprise the bilayered ducts throughout the breast ^18^. We observed that eight of the nine (miR-16-5p not in dataset) miRNAs deregulated during development and BRCA1 mutated tumours were also differentially expressed in these subsets of the human breast (Figure 2B). MiR-485 and miR-542 were enriched in the stem-like population whereas miR-29b-1 and miR-30b were enriched in luminal epithelial cells, when compared to the stem-like and myoepithelial cells, with expression patterns highly similar to BRCA1. MiR-34b, miR-744, miR-485 and miR-542 expression all negatively correlated to BRCA1 expression, which is consistent with our findings from Brca1 knockout glands and human BRCA1 mutant carriers.

### BRCA1 binds to miRNA promoters and regulates miR-29b-1-5p levels in breast cancer

Given the non-canonical role of BRCA1 as a transcriptional regulator we analysed ChIP-seq data for BRCA1 binding in proximity to our differentially expressed miRNAs. BRCA1 binding, was identified for eight of the nine miRNA genes (Table 1). For miR-485-3p, miR-744, miR-664, miR-221, miR-29b-1-5p and miR-30b no histone marks indicative of promoter activity was identified within 15kb upstream of these genes. To more accurately ascertain binding sites that may contribute to the transcriptional regulation of these six miRNAs, in addition to the first 15kb upstream of each gene, the first putative promoter element located upstream of each miRNA was also included in the analysis. These putative promoter elements were enriched in H3K27ac, H3K4me3, H3K4me1 and DNase sensitivity found at promoters. This combined analysis demonstrated that seven of the nine miRNA genes had BRCA1 ChIP-seq binding suggesting a potential mechanism whereby BRCA1 regulates the expression of these miRNAs in breast cancer.

**Table 1.** ChIP-seq and bioinformatic promoter analysis shows that BRCA1 binds to putative miRNA promoter regions. Six of the nine prioritized miRNAs contain BRCA1 ChIP-seq peaks within 15kb of the miRNA transcriptional start site. MiR-744 has a BRCA1 associated ChIP-Seq peak at its nearest H3K4me3 promoter mark. Highlighted in bold are miRNA genes that a greater number of binding sites than is expected by chance.

| hg19 |  |  | 15Kb Upstream |  |  |  | Nearest Promoter Histone Mark |  |  |  |  |
| --- | --- | --- | --- | --- | --- | --- | --- | --- | --- | --- | --- |
| miRNA | Chr | Start | 15KB Upstream | BRCA 1 pAb | BRCA 1 mAb | RNA-Pol II P-Ser2 | Nearest H3K4me3 | Len. | BRCA1 pAb | BRCA1 mAb | RNA-Pol II P-Ser2 |
| miR-34b | 11 | 111383663 | 111368663 | - | 1 | - |  |  |  |  |  |
| miR-485-3p | 14 | 101521756 | 101506756 | - | - | 2 | chr14:<br>101,363,445-101,371,834 | 8389 | - | - | - |
| miR-744 | 17 | 11985216 | 11970216 | - | 1 | 3 | chr17:<br>11,923,129-11,928,868 | 5739 | 1 | - | 2 |
| miR-542-3p | X | 133675467 | 133690467 | 1 | 2 | 3 |  |  |  |  |  |
| miR-664 | 1 | 220373961 | 220388961 | - | 3 | 3 | chr1:<br>220,392,260-220,399,591 | 7331 | - | - | 1 |
| miR-221 | X | 45605694 | 45620694 | - | 2 | 1 | chrX:<br>45,622,880-45,634,966 | 12086 | - | 2 | 1 |
| miR-16-2 | 3 | 160122533 | 160107533 | 1 | 2 | 5 |  |  |  |  |  |
| miR-29b-1 | 7 | 130562298 | 130577298 | - | 2 | 4 | chr7:<br>130,591,395-130,602,861 | 11466 | - | 2 | 2 |
| miR-30b | 8 | 135812850 | 135827850 | 1 | 1 | 3 | Chr 8:135,842,014-135,846,518 | 4504 | 1 | 1 | 1 |

The basal-like breast cancer cell line, HCC1937, harbours a nonsense mutation in the *BRCA1* gene leading to a premature stop codon prior to the DNA binding domain. Using these cells, we stably overexpressed the full-length BRCA1 and assessed expression of the seven miRNAs with BRCA1 binding (Figure 3A). Of these, the expression of miR-29b-1-5p was significantly upregulated, consistent with our Brca1 knockout data and sequencing from the BRCA1 mutant tumours (Figures 3A, 1A and 1B). Six BRCA1 binding sites (enriched from the monoclonal antibody) were identified upstream of the *miR-29b-1* gene, widespread DNase sensitivity is also seen, in conjuction with histone modifications typical of active promoters and enhancers (Figure 3B). Notably, the primary-miRNA-29b encodes both miR-29b-1-5p and miR-29b-3p. MiR-29b-3p expression does not change upon BRCA1 overexpression, suggesting that both transcriptional and post-transcriptional regulation processes are likely involved (Figure 3A).

**Figure 3.**
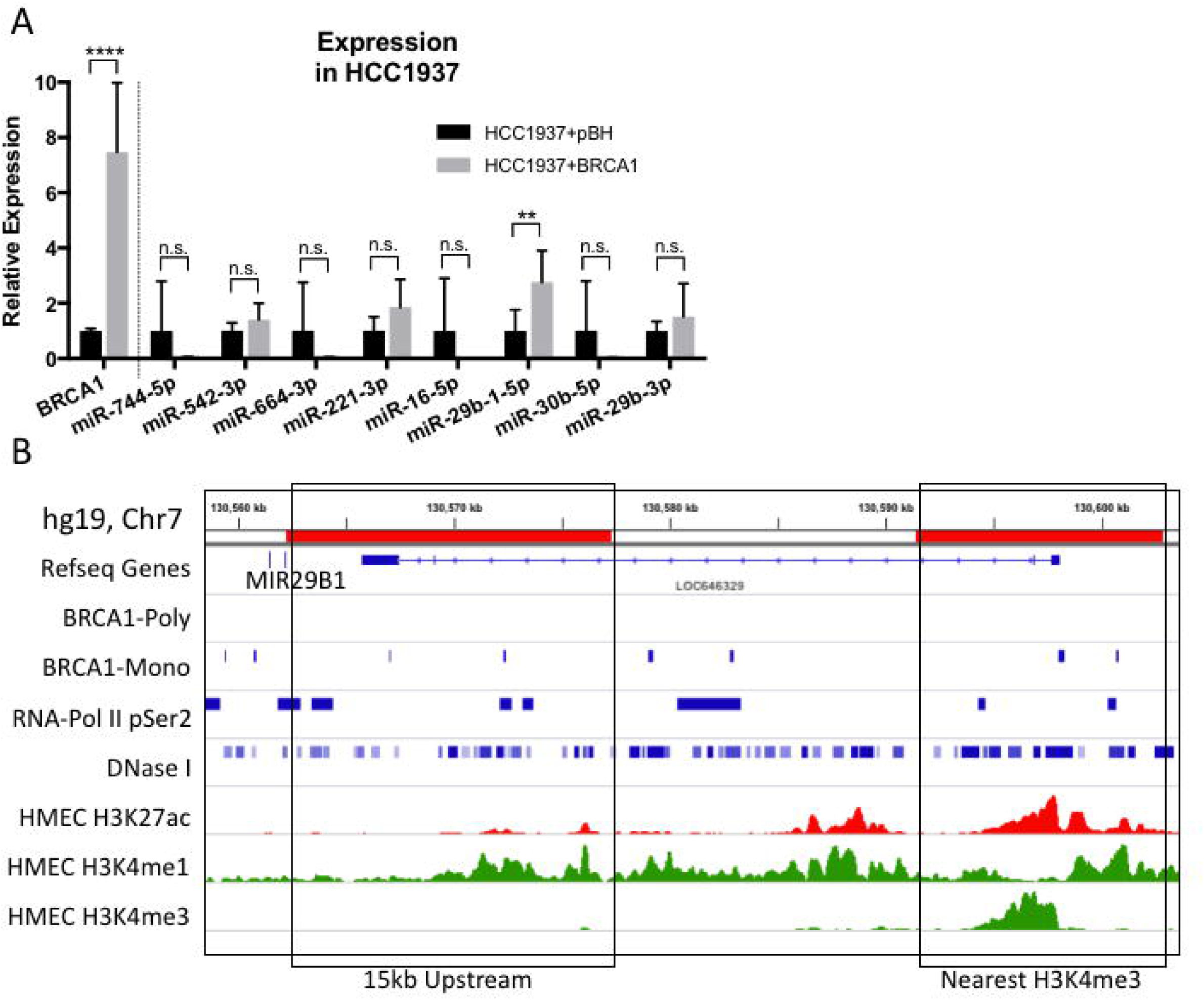
BRCA1 over-expression upregulates miR-29b-1-5p. **A,** relative expression of BRCA1 and seven prioritized miRNAs in the HCC1937 cell line transduced with pBH or pBH-BRCA1 compared to RPLP0 and RNU6B respectively. Results are characteristic of 3 independent replicates. Statistical significance was measured by Students’ t-Test (\*\**P≤0.01*, \*\*\*\**P ≤ 0.0001*, *n.s*.=not significant). **B,** BRCA1 binding proximal to the miR-29b-1-5p genomic loci (miR29a, miR29b). The first 15kb upstream and the nearest H3K4me3 peak are shown, representing the most likely putative promoter regions for this miRNA.

Analysis of possible targets for miR-29b-1-5p reveals that the top 4 targets, USP28, NEUROD1, LIN9 and WDR26 have all been previously associated with breast cancer (Supp Table 3). Interestingly, SPIN1 is among the predicted targets and have been previously identified to be directly targeted by miR-29b-1-5p in breast cancer cell lines^19^.

### Deregulated miRNAs are altered in non-familial breast cancers

BRCA1-associated tumours bear resemblance to TNBC and basal-like tumours, suggesting conserved biological mechanism of tumourigenesis for this tumour types. To determine if these nine miRNAs are also differentially expressed in non-familial breast tumours, including basal-like tumours, expression was analysed in the METABRIC cohort of patients. A heat map of normalised expression demonstrates that these miRNAs are differentially expressed in both familial and non-familial breast tumours, particularly miR-16-2, miR-30b, miR-221 and miR-542-3p (Figure 4).

**Figure 4.**
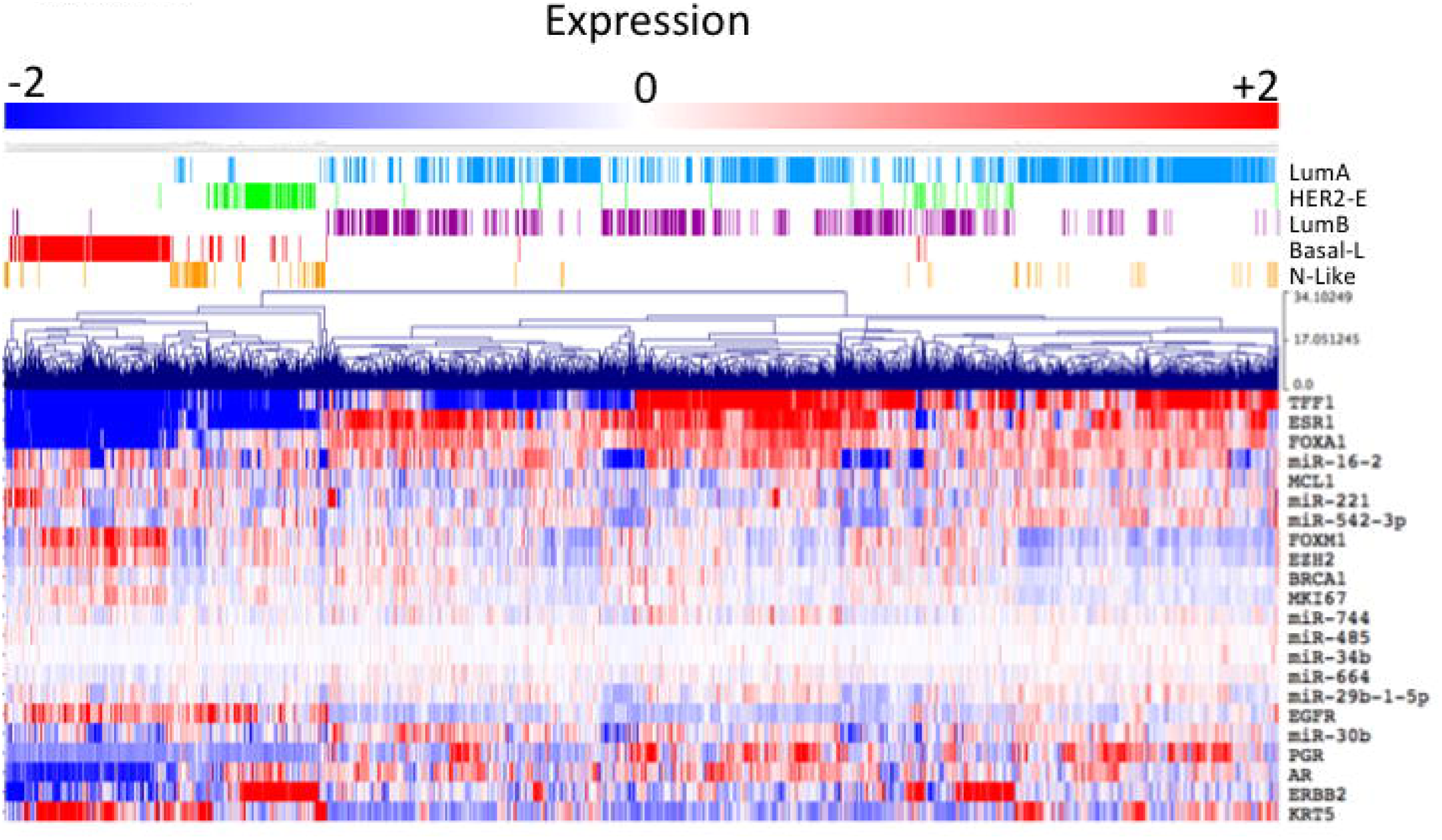
Deregulated miRNAs are also altered in non-familial breast cancers. Deregulated miRNAs display heterogeneous expression in sporadic human breast cancers. Expression data was sourced from the METABRIC cohort with PAM50 subtypes indicated above the heatmap. A number of commonly used molecular markers (*TFF1, ESR1, FOXA1, MCL1, FOXM1, EZH2, MKI67, EGFR, PGR, AR, ERBB2* (*HER2*) and *KRT5*)) of breast cancer subtypes were also included in the heatmap to enable clustering into the PAM50 subtypes.

### BRCA1-associated miRNAs serve as significant biomarkers for survival of basal-like breast cancer

Breast tumours arising in BRCA1 mutation carriers most closely resemble basal-like breast tumours ^3,5^. These tumours do not commonly express the estrogen (ER-) and progesterone receptors (PR-), or HER2 (HER2-). Patients with these tumours have decreased chances of survival and there are currently no targeted therapies available. To determine whether differential expression of the nine miRNAs associated with survival in basal-like tumours, the expression of each miRNA was used to stratify the overall survival (OS) of basal-like breast tumours from the METABRIC cohort. This analysis demonstrated that miR-29b-1-5p greatly stratified survival with high expression associating with a better outcome (HR = 0.281, P-value = 0.0006; Table 2, Supp Figure 1). Three other miRNAs also significantly stratified OS, high miR-664, expression was associated with poor survival whereas high miR-16-2 and miR-744, expression was associated with a better outcome (Table 2). Interestingly, miR-29b-1- 5p stratified OS was greater than any of the tested clinical markers including age at diagnosis and lymph node status. Utilising a multivariate analysis, only five conditions were needed to stratify OS of these basal-like tumours, miR-29b-1-5p, age at diagnosis, lymph node status, ERBB2 and miR-664. These data suggest that BRCA1 deregulated miRNAs can serve as potent biomarkers for the stratification of survival in patients with basal-like breast tumours.

**Table 2.** Prognostic potential of the nine conserved miRNAs in basal-like breast cancer. MiR-29b-1-5p was the most significantly prognostic miRNA entity when interrogated under the univariate and multivariate cox proportional hazard models. HR, CI, ns need explanations.

| Condition | Univariate Cox-proportional hazards model |  |  | Multivariate Cox-proportional hazards model (stepwise) |  |  |
| --- | --- | --- | --- | --- | --- | --- |
|  | HR | (95% CI) | P-Value | HR | (95% CI) | P-Value |
| miR-29b-1-5p (high vs low) | **0.281** | **0.121 - 0.653** | **0.0006** | **0.305** | **0.130 - 0.717** | **0.0067** |
| Age (<40, 41-60, >60) | **0.547** | **0.375 - 0.799** | **0.0018** | **0.557** | **0.365 - 0.848** | **0.0067** |
| Lymph Node (+, -) | **2.317** | **1.340 - 4.007** | **0.0022** | **1.963** | **1.117 - 3.451** | **0.0198** |
| ERBB2 (high vs low) | **3.459** | **1.633 - 7.326** | **0.0052** | **3.875** | **1.799 - 8.346** | **0.0006** |
| miR-664 (high vs low) | **1.983** | **1.069 - 3.679** | **0.0224** | **1.981** | **1.051 - 3.731** | **0.0354** |
| miR-16-2 (high vs low) | **0.542** | **0.313 - 0.941** | **0.0263** | ns |  | ns |
| miR-744 (high vs low) | **0.562** | **0.333 - 0.948** | **0.0322** | ns |  | ns |
| Menopausal Status (Post, Pre) | 1.597 | 0.944 - 2.701 | 0.0811 | ns |  | ns |
| Size (T1, T2, T3) | 1.459 | 0.925 - 2.299 | 0.1072 | ns |  | ns |
| Treatment (Yes, No) | 1.826 | 0.777 - 4.290 | 0.138 | ns |  | ns |
| BRCA1 (high vs low) | 0.653 | 0.366 - 1.165 | 0.14 | ns |  | ns |
| miR-485-3p (high vs low) | 1.508 | 0.863 - 2.637 | 0.1427 | ns |  | ns |
| miR-542-3p (high vs low) | 0.653 | 0.373 - 1.141 | 0.1477 | ns |  | ns |
| miR-221 (high vs low) | 0.686 | 0.370 - 1.272 | 0.2195 | ns |  | ns |
| miR-30b (high vs low) | 1.312 | 0.778 - 2.213 | 0.3132 | ns |  | ns |
| miR-34b (high vs low) | 0.778 | 0.456 - 1.327 | 0.3553 | ns |  | ns |
| Tumour Grade (1,2,3) | 1.443 | 0.600 - 3.468 | 0.3866 | ns |  | ns |
| ESR1 (high vs low) | 0.774 | 0.242 - 2.473 | 0.6545 | ns |  | ns |
| PGR (high vs low) | 1.190 | 0.292 - 4.855 | 0.8141 | ns |  | ns |

## Discussion

Significant molecular and structural alterations occur in the normal breast prior to the development of BRCA1 associated breast cancer. Several studies provide evidence heterozygous loss of BRCA1 contributes to aberrant mammary gland development through increasing DNA damage ^20,21^. MiRNAs have diverse roles in cancer, including effect DNA damage and have previously been linked to BRCA1 associated tumours ^16,22,23^. Advances in therapeutic targeting of miRNAs have made them an attractive target in treating patients with cancer ^24,25^. We present here, that miRNAs deregulated following Brca1 loss may be useful as prognostic markers in patients with basal-like breast cancer or tumours arising in BRCA1 mutant carriers.

We identified differentially expressed miRNAs in Brca1 deficient mammary epitheial cells, human breast tumours and the developing mammary gland. The promoter of miR-29b-1-5p, contains several binding sites for BRCA1 among enrichment of active histone modifications and DNase sensitivity. Rescue of the BRCA1-mutant breast cancer cell line, HCC1937, with overexpression of a full-length BRCA1 gene resulted in a significant increase in expression of miR-29b-1-3p. This miRNA also proved to be a potent biomarker in the progression of basal-like breast cancer, a subtype with significant similarities to BRCA1 associated breast cancer. Indeed, miR-29b-1-3p was shown to be a more significant biomarker for the stratification of overall survival compared to more commonly utilised clinical biomarkers such as lymph-node status and age at diagnosis. Our work compliments a previous study showing that miR-29b-1-3p stratifies overall survival in a small group (n=27) of breast cancer patients with ER- disease ^26^.

BRCA1 is capable of regulating gene expression through direct binding to promoters of protein-coding genes and non-coding RNAs ^10^. For example, wild-type BRCA1 binds to the promoter of miR-155 in breast cancer cells, where it recruits histone deacetylase 2 (HDAC2) to promote chromatin condensation and gene repression REF10. The BRCA1 R1699Q risk variant results in an altered BRCA1 that can no longer recruit HDAC2 and increased miR-155 expression and tumourigenesis. Given BRCA1 also binds upstream of miR-29b-1-5p, it would be interesting to explore the effect of risk variants on BRCA1’s ability to bind and possibly influence miR-29b-1- 3p expression.

The miR-29 family have already been reported to have tumour suppressor activity ^27^. A previous study found many miR-29 family members have complimentarity to DNA methytransferases and that overexpression of miR-29 miRNAs restored normal DNA methylation and decreased tumourigenesis in lung cancer ^28^. Similarly, in breast cancer, miR-29b has been shown to decrease metastasis following stimulation by GATA3, through targeting of regulators of angiogenesis, collagen remodelling and proteolysis ^29^. Recent work has shown that in TNBC cell lines, miR-29b-1-5p acts to suppress oncogenic properties such as viability, apoptosis and migration, and overexpression of the miRNA increased the sensitivity of these cell lines to chemotherapeutic agents^19^. This tumour suppressive role was mediated through direct targetting of the SPIN1 3’ UTR effecting the downstream signaling of WNT/β-catenin and Akt. These data strengthen our finding that high expression of this miRNA is a biomarker for good prognosis. Very little work has explored the role of miR-29b-1-5p in development or cancer, however a single study has found that miR-29b-1-5p and miR-29c can influence proliferation, cell cycle and apoptosis in bladder cancer ^30^. MiR-29b-1-5p is the 5’ miRNA processed from the same stem loop as miR-29b (3’ miRNA), their transcriptional regulation should be identical, but differ in the processing and maturation ^31^. Given that we didn’t detect differential expression of miR-29b in our miRNA-seq data or in BRCA1 over-expressing HCC1937 cells, it is possible that Brca1 regulates the processing of only miR-29b-1-3p through alterations to the miRNA biogenesis and processing pathway ^32^. Future work aiming to understand the role of Brca1 in miR-29b-1-5p regulation should explore possible changes to miRNA processing.

The predicted human targets of miR-29b-1-5p reveal genes previously associated with breast cancer. USP28 is a deubiquitinase that has been shown to stabalise LSD1 and MYC, an important regulator of gene expression, to promote a cancer stem-cell like state and proliferation ^33,34^. Data from Drago-Ferrante *et al* demonstrate that miR-29b-1-5p overexpression suppresses self renewal of mammospheres formed by breast cancer cell lines possibly through miR-29b-1-5p mediated reduction in MYC ^19^. In a seperate study, WDR26 was shown to stablise the Akt signaling complex to promote tumour cell growth and metastasis ^35^. In the Drago-Ferrante paper, they also demonstrated that miR-29b-1-5p is able to repress Akt signalling ^19^, this could possibly be through its putative regulation with of WDR26. MYC is a major oncogene in many cancers, including in BRCA1 breast tumours, where BRCA1 loss leads to overexpression of MYC contributing to tumourigenes ^36^. It is possible that miR-29b-1-5p serves as a redundant repressor of MYC activity through regulation of USP28 and when loss contributes to oncogenesis.

Several studies have investigated the utility of miRNAs in stratifying triple negative breast cancer (TNBC) or basal-like tumours. A 4-miRNA signature has been developed to stratify TNBC, particular those patients receiving chemotherapy, however none of the miRNAs identified in this study are amongst those four ^37^. A separate group identified several miRNA biomarkers of TNBC, of which miR-16 features prominently ^38^. Consistent with our survival data, the authors show that the expression of this miRNA is protective. The low level of overlap between our miRNAs and those already identified as biomarkers for TNBC or basal-like tumours is likely due to the mechanism of discovery. We have utilised developmental signatures to identify miRNAs strongly associated with BRCA1 biology. Dvinge and colleagues support this notion with the finding that miRNAs do have prognostic potential, however when combining miRNA expression signatures with protein coding genes, this potential is far greater ^39^.

We have demonstrated that deletion of Brca1 in the murine mammary gland results in global changes to miRNA expression. Some of these changes in expression are found in BRCA1 mutation carriers, compared to normal breast tissue. We show that BRCA1 influences expression of miR-29b-1-5p in human breast cancer cell lines, possibly via promoter binding and influencing transcriptional regulation. Finally, we show that miR-29b-1-5p is a potent biomarker for the stratification of overall survival in basal-like breast cancer. These data highlight the need to understand normal developmental processing and how they might translate to key findings in tumourigenesis and biomarker discovery.

## Materials and Methods

### miRNA-sequencing

All libraries were prepared using the Ion Torrent RNA-Seq v2 Kits, following the manufacturers recommended protocol. A combination of Ion Torrent and Proton sequencing was used with Ion PGM OT2 200 kits, Ion PGM 200 Sequencing Kit v2, and 318 sequencing chips and Ion PI Template OT2 200 kit, Ion PI Sequencing 200 kit, and P1 chips respectively. All kits were used according to the manufacturers specifications. Trimmed reads were mapped against mm10 with bowtie1 with the following parameters (n 0 -m 1 -l 15 -p 2 -S –max).

### miRNA-sequencing differential gene expression

For differential gene expression determination Partek Genomis Suite v 6.6 and ANOVA were used, and differentially expressed miRNAs were filtered for significance values < 0.05. Filtering for significance identified 145 miRNAs that were differentially expressed. MiRNA expression values were then mean-centred and hierarchically clustered based on average-linkage and Manhattan distance using Multiple Experiment Viewer ^40^.

### Gene expression

Differential expression of miRNAs in BRCA1 tumours was obtained from the Tanic *et al* dataset ^17^. Analysis of miRNA expression in non-familial breast tumours was done using the METABRIC cohort of breast tumours. Gene expression array log2 intensities were mean-centred and hierarchically clustered as above. Expression of miRNA and BRCA1 in epithelial cells of the human breast was obtained from Hirst *et al* and normalised via mean-centring and Pearson’s correlation coefficients to BRCA1 expression shown in the figure ^18^. It is important to note that this dataset doesn’t distinguish 3p and 5p miRNAs. Gene expression across the developmental stages of the murine mammary gland was extracted from GSE15054 and was normliased as above.

### ChIP-seq analysis

BRCA1 and RNA Polymerase II Serine 2 Phosphorylation ChIP-seq data was sourced from GSE45715. This study used two separate BRCA1 anitbodies, a polyclonal (sc-646) and monoclonal (sc-6954) both from Santa Cruz. Data was mapped to hg19 using BOWTIE and ChIP-seq peaks called with MACS14 ^41,42^. For binding associated with miRNA promoter in the MCF10A cell line, BRCA1 binding within 15kb upstream or at the nearest Histone 3 lysine 4 tri-methylation (H3K4me3 mark of promoter elements) was considered relevant. To reduce false positives when looking within the 15kb upstream only binding above background was considered significant. The BRCA1 monoclonal antibody should bind every 15kb within the genome and the polyconal 85kb, based on 203894 and 36655 peaks respectively across the 2.9x10^9^ genome size.

### Tissue culture

The HCC1937 breast cancer cell line was grown in RPMI-1640 supplemented with 10% Fetal Bovine Serum, 1 mM Sodium Pyruvate, 1:100 Penicillin/Streptomycin (Thermo Fisher Scientific) and 10 mM HEPES (Sigma-Aldrich). HCC1937+BRCA1 were sourced from Deans and McArthur 2004^43^.

### qRT-PCR

RNA was extracted from HCC1937 cells using Trizol and phenol-chloroform extraction. cDNA was produced using Superscript III following the manufacturers protocol. qRT-PCR was performed with the Qiagen miScript SYBR based system. Primers for miRNA expression were, mm-miR-206-3p, hsa-miR-30b-5p, hsa-miR-744-5p, hsa-miR-664-3p, hsa-miR-16-5p, hsa-miR-485-3p, hsa-miR-34b-5p, hsa-miR-542-3p, hsa-miR-29b-1-5p, hsa-miR-29b-3p, hsa-miR-221, and RNU6b, sequences and catalogue numbers listed in Supp Table 4. Human miRNA primers were used when mouse were not available, miRNAs sequence is conserved between species. BRCA1 expression was performed using Taqman reagents, following the manufacturers protocol, probes BRCA1 (HS01556193_M1) and RPLP0 (HS99999902_M1).

### Tumour survival analysis

Expression and clinical annotations were sourced from METABRIC ^39,44^. MedCalc was used to analyse association of miRNA expression with clinical outcome. For assignment of high and low expression groups, receiver operator curves (ROC) were employed to find the optimal expression cutoff for survival analysis. Clinical markers were divided into groups as indicated in Table 2, size (T1 = ≤20mm, T2 = 20-50mm, T3 = ≥50mm). Univariate Cox-proportional hazards ratios were determined in MedCalc, following by a Multivariate Cox-proportional hazard test using a stepwise model.

### miRNA Target Prediction

The mmmRNA^45^ database was used to predict human mRNA targets of miR-29b-1-5p. This database combines predictions from four databases on experimental validation of miRNA targets and nine databases on predicted targets. Supp Table 3 contains the full list of targets produced by this database.

## Acknowledgments

This study makes use of data generated by the Molecular Taxonomy of Breast Cancer International Consortium. Funding for the project was provided by Cancer Research UK and the British Columbia Cancer Agency Branch (38).

## Author Contributions

M.J.G.M performed all bioinformatic analysis, including expression and survival analysis, drafted the manuscript and co-designed the study. G.K.S performed all qRT-PCR analysis, drafted the manuscript and co-designed the study. A.W. co-designed the study and contributed to drafting the manuscript. D.K. performed the miR-Sequencing and analyses. B.L.B, A.S, S.L.E and J.D.F contributed to the study design and manuscript drafting. M.A.B conceived the study, designed experiments and contributed the drafting of the manuscript.

## Data Availability

miRNA-Seq from Brca1 condition knockout mammary glands is available upon request.

---

**Supp Figure S1.**
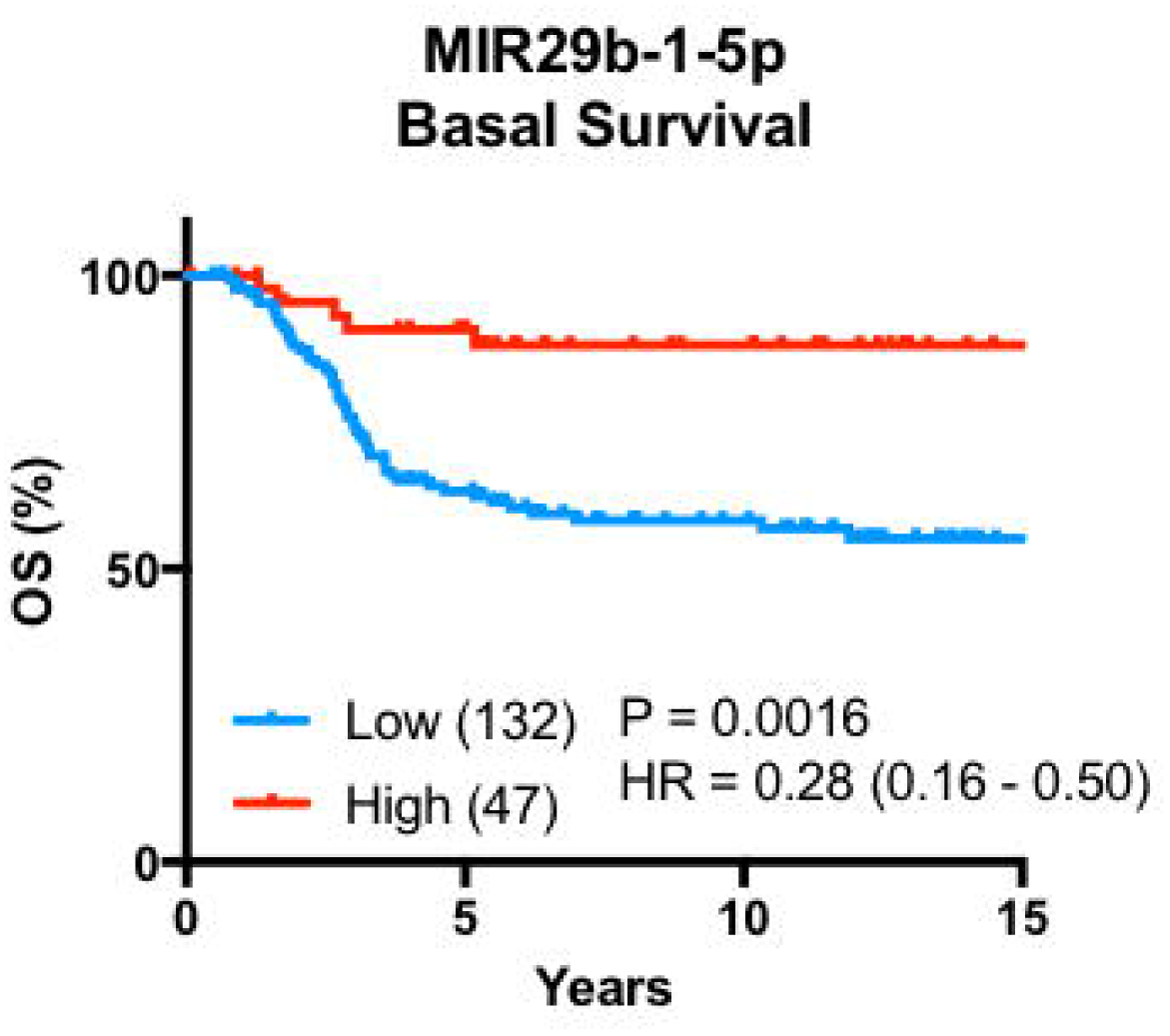
MIR29b-1-5p stratifies overall survival of basal-like breast tumours.

